# Seasonal connectivity of microbes and carbohydrates between ocean, atmosphere, and cryosphere in Kongsfjorden (Svalbard, Arctic Ocean)

**DOI:** 10.64898/2025.12.01.691664

**Authors:** Matthias Wietz, Manuela van Pinxteren, Heike M. Freese, Cathrin Spröer, Sebastian Zeppenfeld

## Abstract

The coupling between ocean and atmosphere across the strong seasonal gradients in the Arctic is poorly understood. Here, we explored the microbial and glycobiological connectivity between the sea surface microlayer (SML), the underlying seawater (ULW), snow, and aerosol particles in Kongsfjorden (Svalbard, 79°N) during autumn and spring. The marked overlap between marine and atmospheric microbiomes illustrates considerable sea-air transfer, linked to seasonally distinct environmental communities. For instance, *Polaribacter* and *Formosa* were aerosolized during the spring bloom, compared to *Colwellia* in autumn. Air-mass trajectories and microbial source tracking revealed a greater marine contribution in autumn, whereas spring aerosols were shaped by stronger winds and the cryosphere. Aerosol particles nonetheless contained numerous unique taxa, including Actinobacteria likely originating from terrestrial sources. Linking bacterial, microeukaryotic, carbohydrate, and meteorological dynamics established an overarching perspective across seasons and habitats, identifying four distinct ecosystem states. Genome-sequenced bacterial model isolates, representing key environmental populations, encode adaptive traits such as carotenoid and ectoine biosynthesis, supporting survival in the SML and atmospheric transfer. Comparison with time-series records from the nearby Fram Strait revealed that many aerosolized bacteria are consistent microbiome components; with implications for ecology and biogeochemistry across the wider Arctic.

## INTRODUCTION

Around the Svalbard archipelago in the high Arctic, microbial and biogeochemical regimes show distinct seasonality [1–4]. Comparable patterns occur in the Arctic atmosphere [5–7]. However, the seasonal connectivity between aquatic and atmospheric realms – integrating microbiological, chemical and ecological dynamics – remains underexplored.

Aerosol particles are key connectors between seawater and atmosphere, mediating the transfer of diverse microbial and chemical components [8–10]. The aerosolization of waterborne components mainly occurs through bubble bursting at the surface microlayer (SML). Organic matter and microbial cells can be enriched up to 1000 times in the SML compared to the underlying water (ULW), coincident with specific chemical signatures and microbial adaptations [11–14]. The aerosolization of microbes aligns with their genetic and phenotypic traits [15, 16], and can vary by season and the environmental origin; for instance terrestrial, marine, or cryosphere [17–19] Meteorological conditions play a key role in this context [20–22], including atmospheric dispersal over considerable distances [23–26].

The strong seasonality in the Arctic – featuring seasonally distinct microbial taxa, genetic functions, and substrate regimes – likely coincides with marked contrasts in ocean-atmosphere connectivity. Arctic aerosol particles mostly originate from local marine sources, especially the SML, plus long-range advection and terrestrial input [5, 27, 28]. These dynamics align with the phytoplankton bloom stage and associated production of carbohydrates, amino acids, and volatiles [29]. For instance, D- and L-amino acids have been suggested as “aerosol indicators” for early- and post-bloom respectively; together with the enrichment of specific microbes along the bloom succession [28, 30]. Notably, aerosolized compounds might undergo microbial transformation in the air [21], supported by laboratory experiments [31].

Here, we explore microbial communities and carbohydrates in the SML, the corresponding ULW, and aerosol particles during autumn 2021 and spring 2022 in Kongsfjorden, Svalbard (79°N). These times mark the early and late phytoplankton bloom, featuring distinct microbial genera and environmental conditions [2]. Kongsfjorden and adjacent waters have been extensively studied regarding microbial, biogeochemical and environmental dynamics across seasons and years [32–38]. We hypothesize that seasonal changes in marine-ecological and meteorological conditions shape both the waterborne and atmospheric microbiome; from autotrophic phytoplankton to heterotrophic bacteria. Carbohydrates are a key component of these dynamics [39, 40]. Accordingly, a study conducted alongside the present work has revealed pronounced seasonality in marine carbohydrate budgets and their atmospheric transfer [41]. We demonstrate marked sea–air connectivity of prokaryotes, microeukaryotes and carbohydrates, supported by genomic analyses of model isolates and seasonal microbiome records from the nearby Fram Strait [42, 43]. Given the exceptional warming of the Arctic [26] and the environmental impact on the airborne microbiome [44], the sources and sinks of microbes and molecules are key aspects for understanding the microbial loop.

## RESULTS AND DISCUSSION

Environmental samples were collected in autumn 2021 and spring 2022 in Kongsfjorden, Svalbard (79°N 11°E; Supplementary Tables S1-S3). Aerosol particles were collected over 4–7 day intervals next to the shoreline, with parallel SML and ULW sampling across Kongsfjorden. Snow was sampled in spring 2022, around 500 m inland. Microbiome dynamics were contextualized with carbohydrate concentrations, meteorological conditions, genomic traits of model isolates, and year-round ecosystem records from the nearby Fram Strait [42, 43].

### Overall microbiome structure

SML und ULW microbiomes were largely congruent; while aerosol particles harbored distinct communities with greater internal variability (Figure 1). Nonetheless, multiple taxa were shared across all niches (see below). All communities showed a clear seasonal signal (PERMANOVA; *p* < 0.001). Alpha-diversity was significantly higher in autumn across sample types (Dunn’s test with Benjamini-Hochberg correction; *p* < 0.05; Figure 1). In contrast, bacterial diversity in the adjacent Fram Strait peaks in April [2], likely corresponding to different oceanographic and stratification conditions. SML and ULW featured significantly higher richness than aerosol particles, whereas autumn aerosol particles featured higher diversity and evenness (Dunn’s test with Benjamini-Hochberg correction; *p* < 0.05).

**FIG 1:**
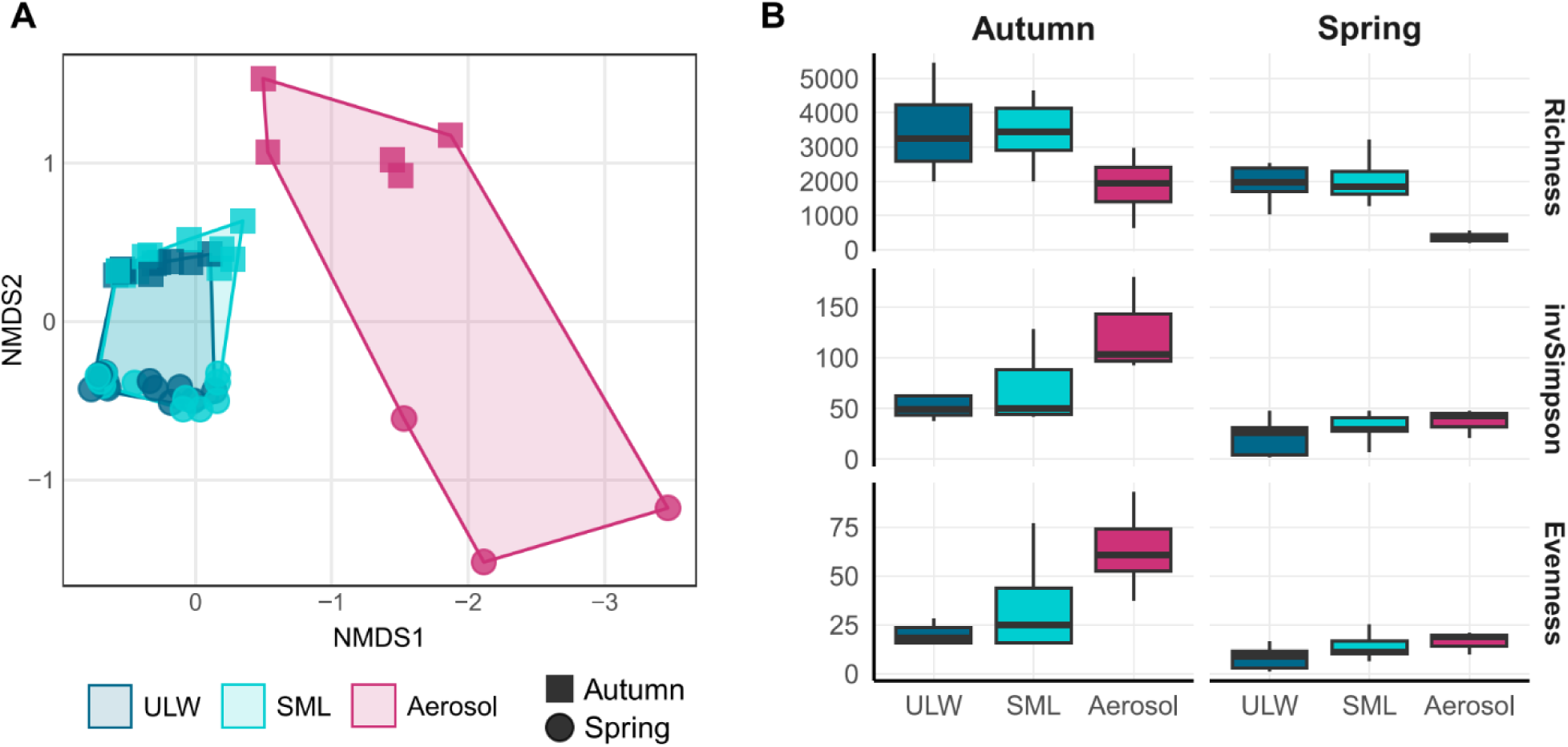
Seasonal microbiome diversity. Non-metric multidimensional scaling (stress value 0.07) and alpha-diversity of bacterial communities in the surface microlayer (SML), the underlying seawater (ULW), and aerosol particles.

Gammaproteobacteria showed comparable abundances across seasons and sample types (Figure 2). In contrast, Alphaproteobacteria and Bacteroidetes peaked in autumn and spring respectively; comparable to the Fram Strait [2]. The prevalence of Verrucomicrobia in autumn SML and ULW underlines previous reports from Kongsfjorden [45]. Aerosol particles harbored more Actinobacteria and Planctomycetes (Figure 2), likely linked to non-marine sourcing (see below). Among SML and ULW communities, sequential filtration allowed distinguishing microbial size classes (planktonic bacteria vs. attached bacteria / larger eukaryotes). However, season was a stronger delineator of community variability (explaining 14% vs fraction explaining 6%; PERMANOVA, *p* < 0.001).

**FIG 2:**
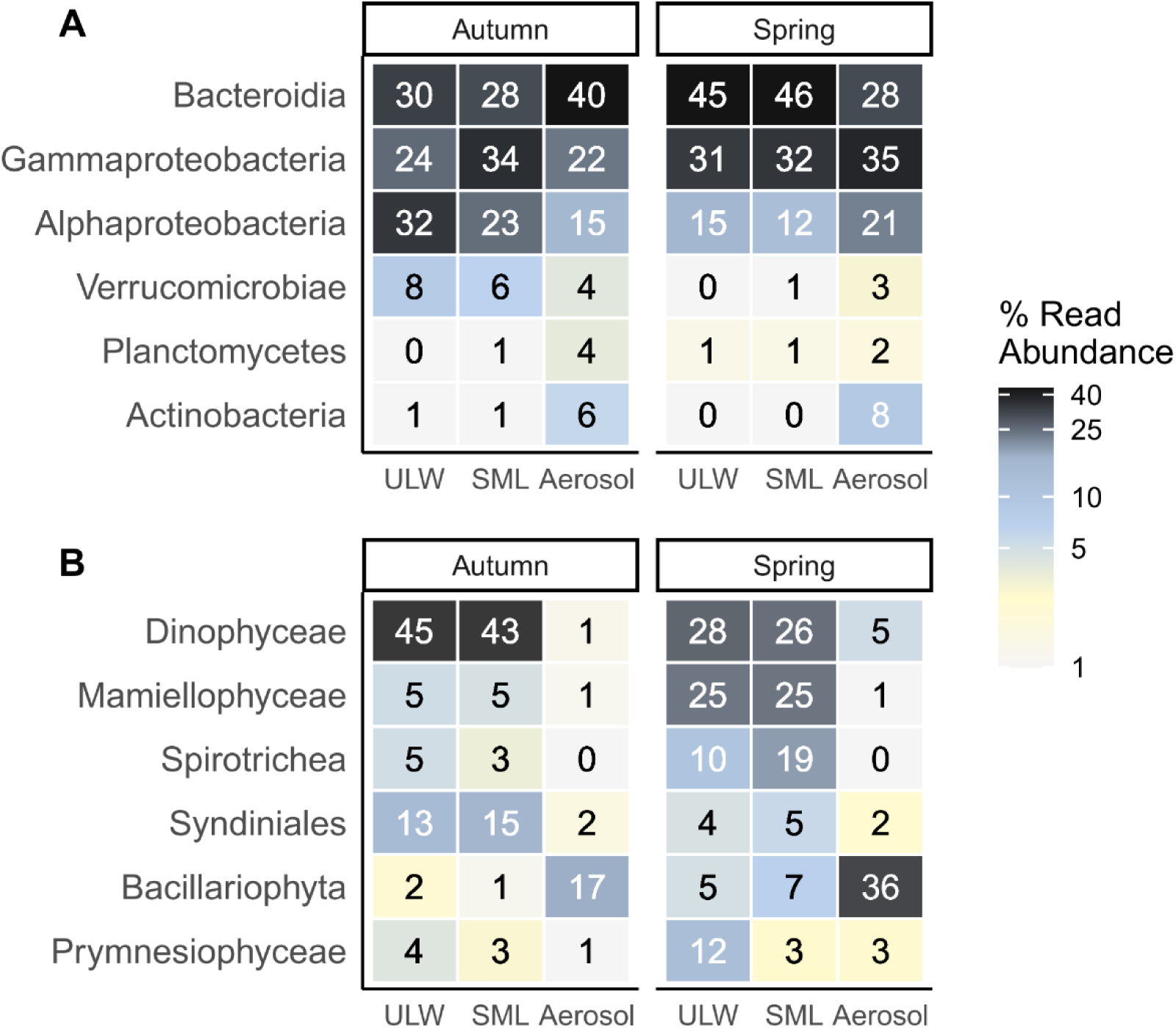
Seasonal microbiome composition. Average relative abundances of the major bacterial and microeukaryotic genera in SML, ULW, and aerosol particles.

Microeukaryotic communities showed comparable seasonality (Figure 2). The predominance of Dinophyceae and Syndiniales in autumn reflects mixo- and heterotrophy in the late productive season, whereas photosynthetic taxa (prymnesiophytes and diatoms) and flagellates (*Mamiellophyceae* and *Spirotrichea*) predominated in spring. We establish more detailed linkages between trophic levels and their environmental drivers below, but subsequently focus on the aerosolization of bacteria, since larger-sized eukaryotes − constituting lower cell numbers than bacteria − are presumably aerosolized more randomly, resulting in greater stochasticity.

### Microbial distribution and adaptations across niches

We then explored the distribution and connectivity of ASVs across niches (Figure 3). Aerosol particles harbored ∼1300 unique ASVs, which however only constituted ∼8% of the aerosol particle community (i.e. many but rare taxa in line with alpha-diversity; Figure 1). The prevalence of Actinobacteria (Figure 2) corresponded to diverse taxa including *Arthrobacter*, *Nocardioides*, and *Mycobacterium* (Figure 3). Microbial source tracking – i.e. calculating community overlap between niches – showed that aerosol microbiomes were more similar to SML than ULW microbiomes (Figure 3), highlighting the SML’s role in sea-air connectivity [46]. Still, the majority of aerosolized ASVs occurred in both SML and ULW, constituting an average abundance of ∼58%. This considerable microbial connectivity between ocean and atmosphere included *Polaribacter*, *Formosa* and *Methylophagaceae* (Figure 3) – key drivers of carbohydrate and methanol cycling during phytoplankton blooms [39, 47]. *Polaribacter* was similarly abundant in aerosol particles during a preceding spring bloom in Kongsfjorden [30], indicating frequent aerosolization during high primary production. *Formosa* correlated with ice-nucleating particle concentrations near Greenland while being metabolically active [28], linking bacterial aerosolization to carbohydrate transformations.

**FIG 3:**
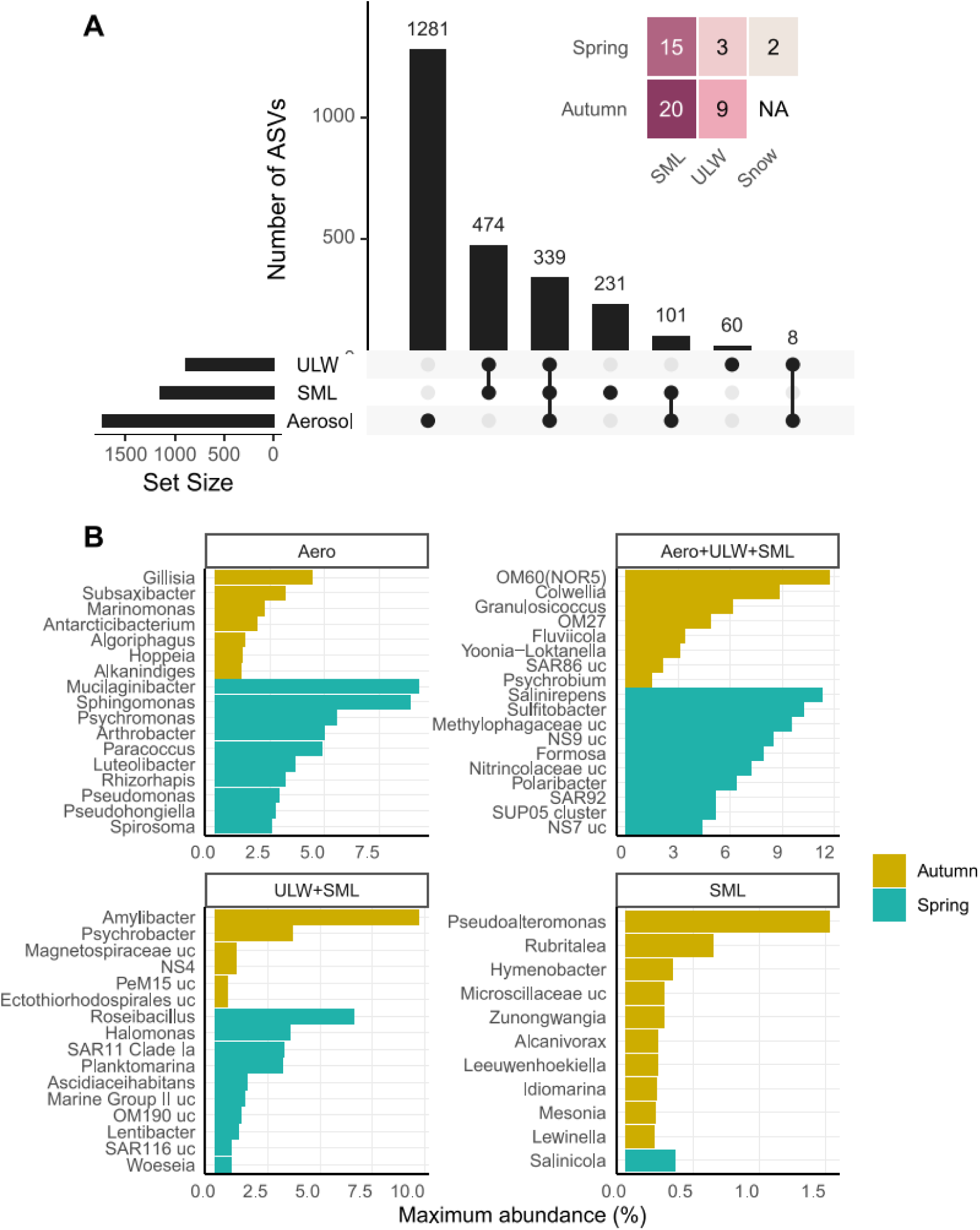
Bacterial composition across niches and seasons. **A:** Shared and unique ASVs in SML, ULW, and aerosol particles. The insert shows % overlap between ULW, SML and snow microbiomes with those in aerosol particles, as determined by source tracking. Autumn was almost snow-free and no snow was sampled; hence no overlap could be calculated. **B:** Predominant genera per niche based on the top50 ASVs, showing the maximum abundance in their peak season.

To quantify niche preferences, we calculated three enrichment indices: aerosol particles vs seawater (Aerosolization Index), ULW vs SML (SML Index), and planktonic vs attached fractions (Attachment Index). SML and Attachment indices peaked in autumn and were positively correlated (Spearman’s *rho* = 0.52; *p* < 0.001), illustrating two key patterns: more particles in the SML, reflecting the known enrichment of organic matter [48], and more PA bacteria in autumn. Accordingly, SML taxa predominated in the PA fraction (∼64%), compared to ULW taxa in the FL fraction (∼63%). The positive correlation between Aerosolization and Attachment indices (Spearman’s *rho* = 0.25; *p* < 0.001) underlines that PA bacteria are favorably aerosolized. Consequently, the mostly free-living SAR11 and *Amylibacter* [49, 50] were absent in aerosol particles. Nonetheless, their presence in autumn fog water collected hundreds of meters inland (Supplementary Figure S2) indicates the presence of atmospheric niches for bacteria with reportedly little aerosolization [15]. The fog water furthermore comprised terrestrial taxa like *Pseudomonas* and *Arcicella*, indicating sourcing from soils.

Adaptations to salinity and UV stress are essential for survival in the SML [51, 52], and our genome-sequenced isolates illustrate the underlying traits. *Salinicola* sp. NYA28a and *Halomonas* sp. NYA30 — matching asv647 and asv310 and hence representing relevant environmental populations — predominate in the SML (Figure 3, Supplementary Figure S1). Both isolates belong to the salinity-tolerant *Halomonadaceae* family and encode identical ectoine biosynthesis clusters (Supplementary Table S3), reflecting adaptation to the rapid osmotic shifts in the SML [51]. NYA28a further encodes an unusual zeaxanthin cluster (*crtE*–*idi*–*crtXYIBZ*) yielding fivefold more carotenoid [53] plus photolyase and lipopolysaccharide biosynthesis genes; together promoting UV resilience [54–57]. The SML is enriched in polysaccharides (Supplementary Figure S3; Benjamini-Hochberg corrected *p* < 0.01), and NYA28a encodes diverse carbohydrate-active enzymes to utilize these: eight GH13 subfamilies, a PL5 / PL14 / PL38 lyase cluster (likely targeting alginate), and a PL22 lyase plus PQQ operon (likely targeting pectin). Together with the considerable sea-air transfer of carbohydrates — measured up to a kilometer via a tethered balloon in parallel to our microbial study [41] — our observations emphasize the connectivity between aquatic and atmospheric niches.

### Meteorological impact and environmental sourcing

Air temperature, pressure, wind speed, and the predominant wind direction differed between autumn and spring (Supplementary Figure S4; Wilcoxon rank-sum test, *p* < 0.05), together contributing to the seasonality of aerosol microbiomes (PERMANOVA, *p* < 0.05). Atmospheric back-trajectories highlighted seasonal contrasts (Supplementary Figure S5), with air flowing across distinct surfaces before reaching Kongsfjorden: sea-ice in spring, open waters in autumn (Figure 4a; Dunn’s test with Benjamini-Hochberg correction, *p* < 0.001). Accordingly, predicting the origin of ASVs shows a higher cryosphere contribution in spring, and higher marine contribution in autumn (Figure 4b; Wilcoxon rank-sum test, *p* < 0.001). Our spring sampling occurred under heavy snow cover plus fresh precipitation from two storms; promoting the occurrence of cryosphere taxa in snow and aerosol particles, such as *Arthrobacter* and *Ramlibacter* (Figure 4c) [58–60]. These dynamics resemble the cryosphere sourcing of spring air microbiomes in Greenland, driven by strong winds [61–63]. The aerosolization of *Tychonema* and *Massilia* reflects their known dispersal through snowstorms [63] and stress resistance in snow [64]. Detection of the lichen symbiont *Rhizorhapis* [65] and permafrost taxa like Gitt-136 [66] indicates additional dynamics during cryosphere-related aerosolization.

**FIG 4:**
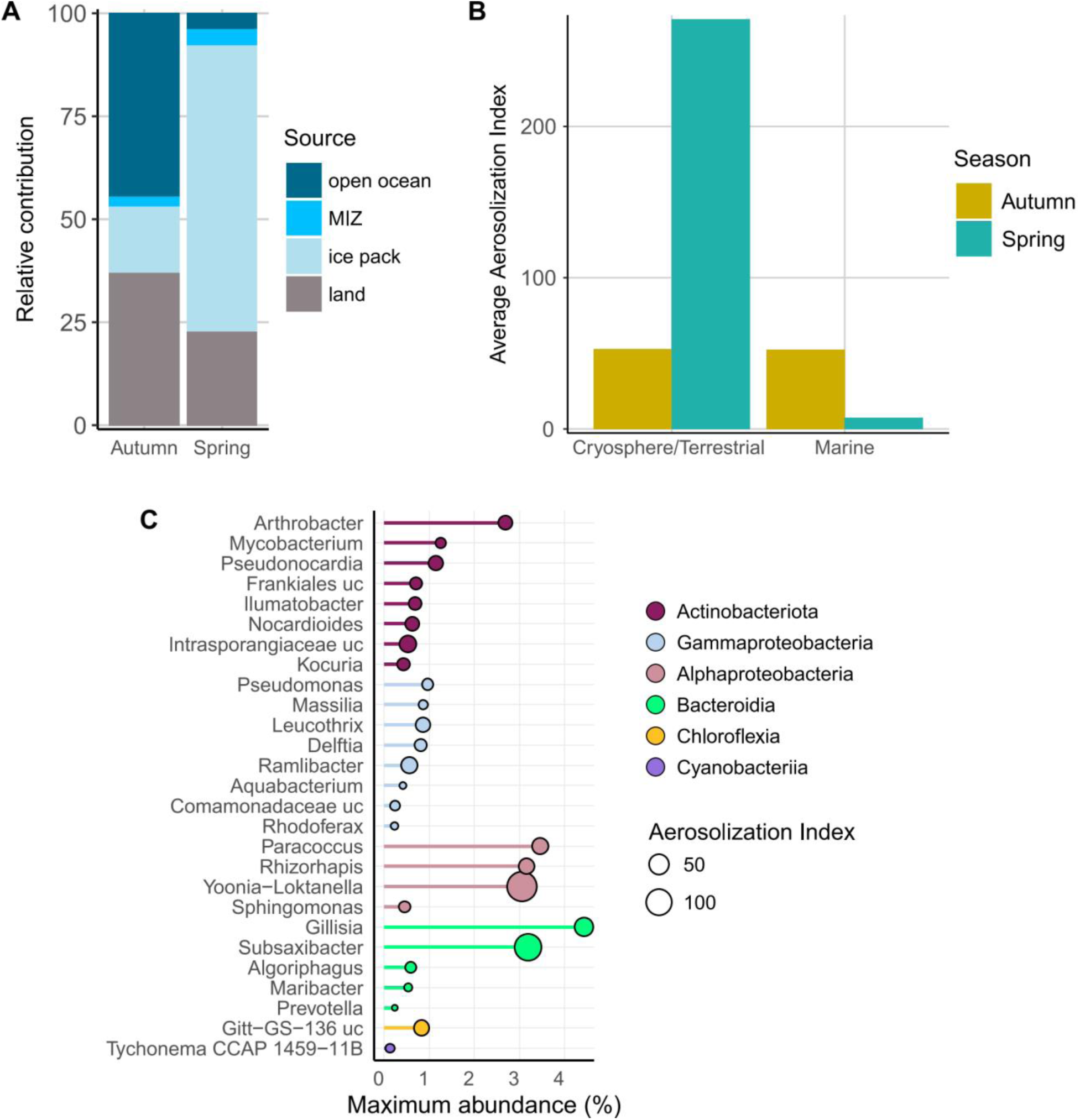
Environmental sourcing and the role of snow in aerosol particle dynamics. **A:** Seasonal trajectories of air masses, showing the environment they predominantly flowed across in the 48 hours before arrival in Kongsfjorden. **B:** Seasonal aerosolization of taxa originating from marine vs cryosphere/terrestrial sources, based on ENVO classifications. **C:** Maximum abundance of bacterial genera co-detected in snow and aerosol particles. Aerosolization Index and taxonomy are illustrated by dot size and color, respectively.

### Seasonal ecosystem context

We contextualized microbial, biochemical and atmospheric seasonality via partial least squares regression (sPLS), identifying four clusters with distinct environmental, temporal, and taxonomic signatures (Figure 5).

**FIG 5:**
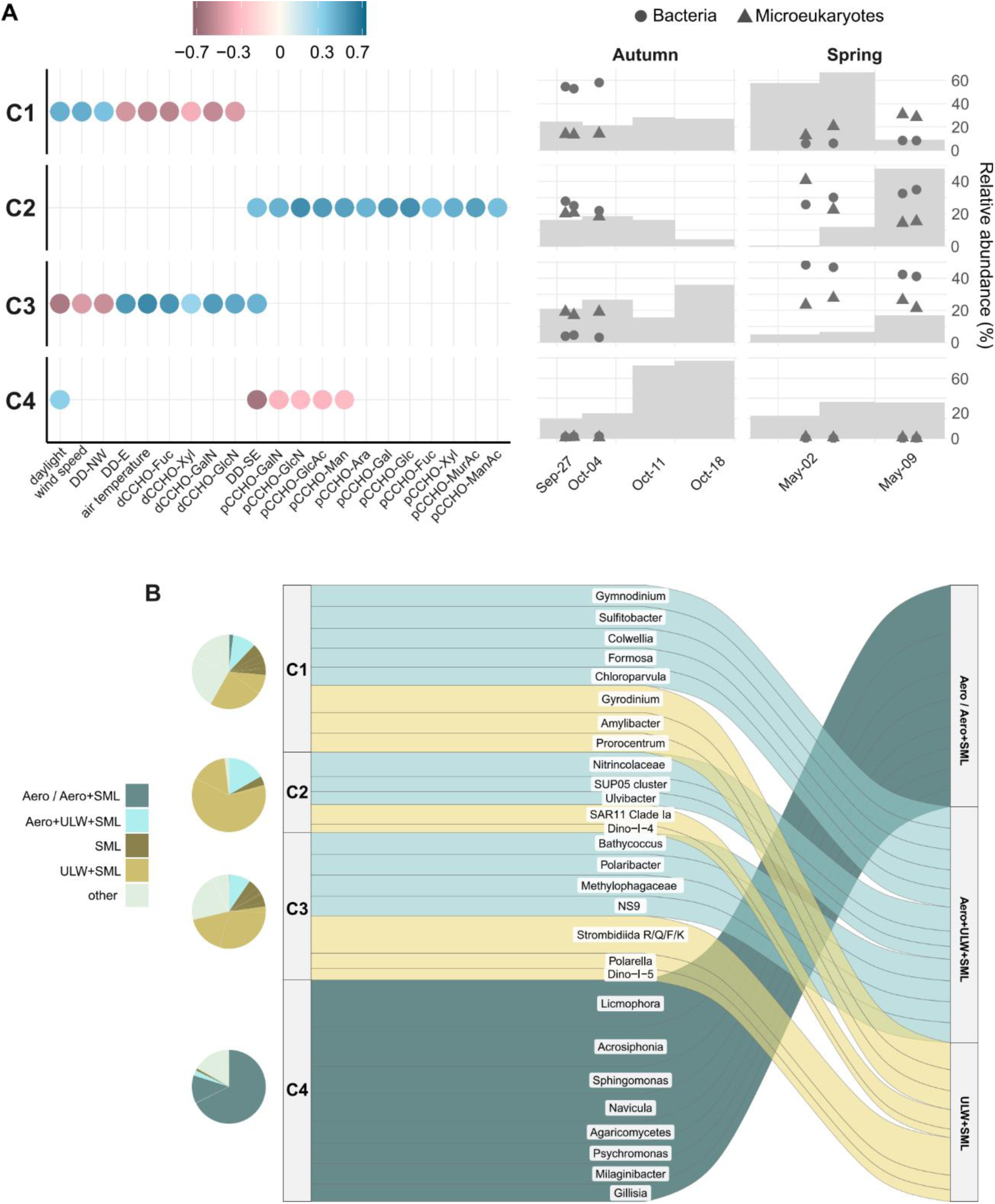
Connections between microbiomes, carbohydrates, and meteorology. **A:** Four microbial clusters with distinct environmental correlations (>|0.5|; left) and temporal dynamics (right). Relative abundances for bacteria and eukaryotes are calculated separately per sample type. dCCHO: dissolved combined carbohydrates; pCCHO: particulate combined carbohydrates; dFCHO: dissolved free carbohydrates. **B:** Contribution of niches to clusters (circles) and major corresponding genera (alluvial plot of square root-transformed abundances).

The autumn cluster C1, associated with dissolved combined carbohydrates, temperature and easterly winds, comprised typical late-bloom microbes, for instance *Gymnodinium* and *Prorocentrum* dinoflagellates [67]. The co-occurring *Amylibacter* and *Sulfitobacter* can degrade the algal senescence compound DMSP [42, 68]. Cluster C1 also harbors *Salinibacterium* asv4472, predominant in SML and aerosol particles (Supplementary Figure S1). The corresponding genome (NYA9b) encodes a flexixanthin-related carotenoid and several other natural products (Supplementary Table S3), indicating substantial competitiveness. Cluster C2 contains taxa with pan-Arctic occurrence across seasons: *Nitrincolaceae*, *Thioglobaceae* and SAR11 Clade Ia, the most abundant genus in ULW+SML. *Polaribacter* and *Bathycoccus*, signature microbes of the spring cluster C3, are indicative of high primary production. The co-occurrence of different Strombidiida clades indicates diverse traits and niches of these ciliates; from autotrophic, heterotrophic to saprotrophic [66–68]. Cluster C4 comprised aerosolized microbes and correlated with southeasterly winds, suggesting regional atmospheric sourcing. The presence of *Navicula*, *Acrosiphonia* and *Licmophora* ASVs, alongside elevated particulate combined carbohydrates, illustrates a contribution of diverse algal taxa and substrates [72–75]. C4 also harbors *Psychrobacter* asv515 with predominance in autumn aerosols and spring snow (Supplementary Figure S1). The corresponding genome (NYA8) encodes a siderophore for iron acquisition and the machinery for assimilatory nitrate reduction to generate biomass from nitrate (Supplementary Table S3). NYA8 furthermore encodes a nitronate monooxygenase with 84% amino acid identity to a functionally characterized homologue from Antarctic *Psychrobacter* [76], suggesting that nitronate metabolism is common at both poles.

### Biogeographic and biogeochemical implications

Multiple Kongsfjorden taxa are shared with microbiomes in the West Spitsbergen and East Greenland Currents (Figure 6), derived from multiannual records from the FRAM Observatory [42, 43]*. Thioglobus* asv5 – occurring in seawater, aerosol particles, and snow – is a “pan-Arctic resident” [42], highlighting sea–air connectivity of key taxa from Svalbard fjords to the central Arctic Ocean. *Thioglobus* asv5 might perform mixotrophic carbon assimilation, based on an isolate from the Yermak Plateau north of Svalbard with 100% rRNA gene identity [77]. *Nitrincolaceae* asv2 from cluster C4 occurs in aerosols and the Atlantic-influenced West Spitsbergen Current, having 100% rRNA identity with isolates from the newly classified, ecologically diverse *Njordibacter* family [77]. *Sphingorhabus* asv19 prevailed in Kongsfjorden snow and the ice-covered East Greenland Current, highlighting the cryosphere contribution in spring including potential long-range advection. The corresponding genome (NYA22) encodes multiple biosynthetic gene clusters, including a zeaxanthin-related carotenoid for UV protection with homologs in Greenland glacier bacteria [54, 78]. Seven ribosomal peptides, a homoserine lactone and a polyketide (Supplementary Table S3) suggest considerable antagonistic and interactive potential. Several genes indicate participation in both nitrification and denitrification (encoding *narGHI* and *norB*).

**FIG 6:**
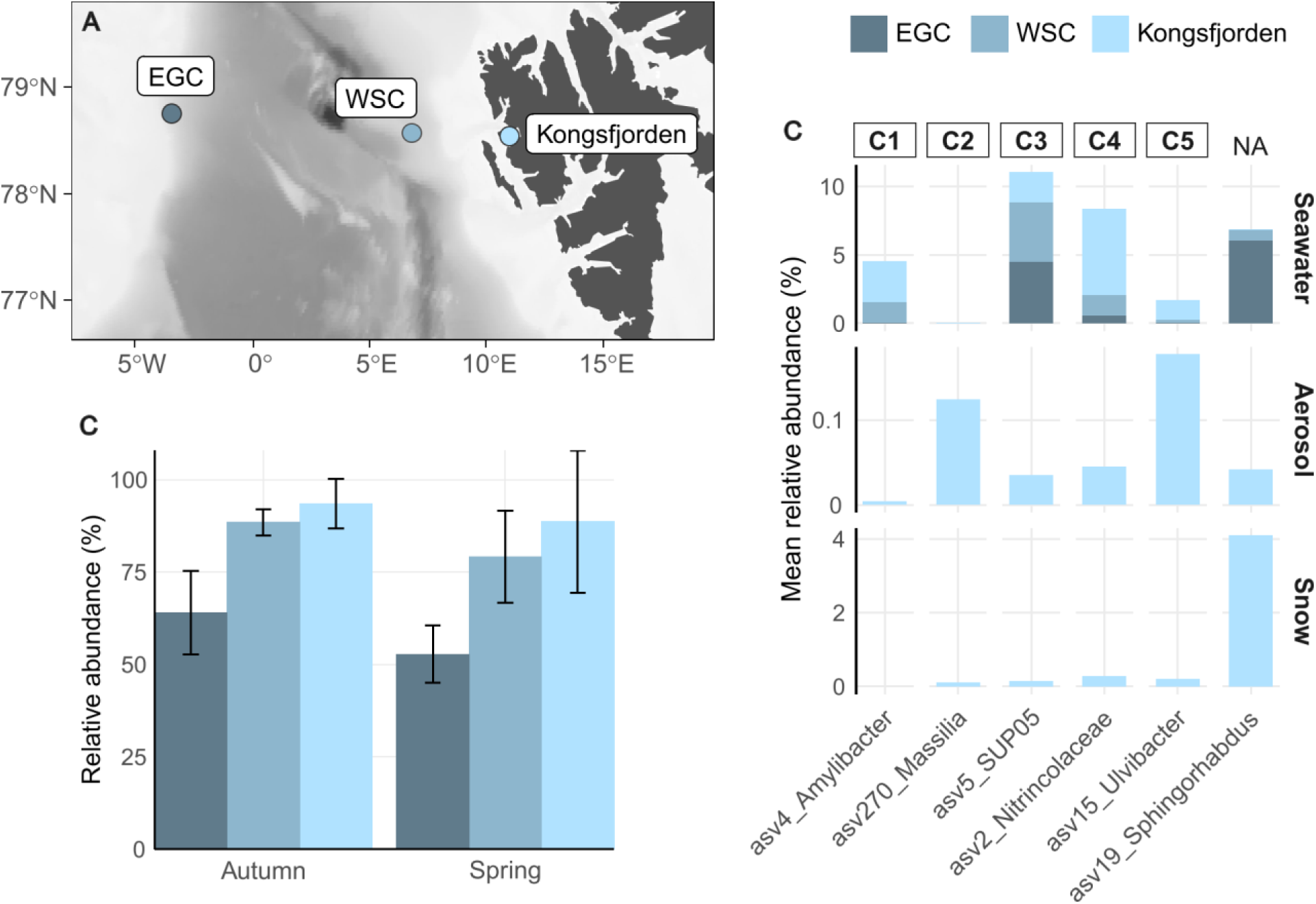
Implications for the wider Arctic. ASV distribution in Kongsfjorden seawater, aerosol particles, and snow in comparison with four-year records from the Fram Strait. **A:** Location of Kongsfjorden and moored autonomous samplers in the Atlantic-influenced West Spitsbergen Current (WSC) and the polar-influenced East Greenland Current (EGC). **B:** Combined relative abundance of Kongsfjorden “core taxa” (occurring in aerosol particles, ULW and SML at mean relative abundance >0.005) by season in comparison to WSC and EGC. **C:** Mean relative abundances of signature ASVs across sites and niches.

## CONCLUSIONS

Our study illustrates how biology, meteorology, and physicochemistry shape the seasonal sea-air connectivity in Kongsfjorden, with broader implications for Arctic microbial ecology and glycobiology. We present three major conclusions:

1. Seasonally distinct “source microbiomes” in context of environmental gradients are a major driver of aerosol particle dynamics; likely promoted by genetic and phenotypic adaptations of seasonal communities [10, 16, 42, 43].
2. Aerosol particle microbiomes scale with meteorological conditions and air mass history, including the transfer of cryosphere microbes in spring. Potentially, aerosolized microbes represent “seeds” when precipitating from the atmosphere, especially when dispersed in dormant stages [23, 79].
3. Microbial similarities between Kongsfjorden, Fram Strait and the central Arctic Ocean indicates pan-Arctic patterns of aerosolization, linked to microbial connectivity via the transpolar drift, Atlantic water influx, and atmospheric dispersal.

## METHODS

### Seasonal sampling

Samples were obtained during the BELUGA campaign at the AWIPEV research station (Ny-Ålesund, Svalbard) in September-October 2021 and April-May 2022 (Supplementary Table S1). Aerosol particles were sampled next to the Old Pier over 4- to 7d intervals on 47 mm polycarbonate membranes, as detailed in [41]. During these intervals, the SML and ULW (1 m depth) were repeatedly sampled from a motorized dinghy [80]. Water was sequentially filtered onto 47 mm polycarbonate membranes; first through 3 µm pore size and the flow-through on 0.2 µm using mild vacuum (200 mbar). All filters were immediately frozen at −20°C until further processing.

### Microbial metabarcoding

Microbial communities were analyzed in line with the FRAM Observatory [2]. Briefly, 16S and 18S rRNA gene fragments were MiSeq-sequenced using primers 515F–926R and 528iF–964iR [81, 82], processed into amplicon sequence variants using DADA2 v1.14.1 [83], and taxonomically classified using Silva v138 and PR^2^ v4.12 [84, 85]. Singletons, mitochondria, chloroplast, and metazoan reads were discarded. To account for potential contaminants, counts from PCR negative controls were subtracted from the raw abundance tables. Sequence results were further carefully inspected manually, and potential contaminants removed by comparison with literature data.

### Bacterial isolation, genome sequencing, and comparative genomics

Bacterial strains were isolated from enrichment cultures in minimal seawater medium [86] with arabinogalactan, fucoidan, or alpha-mannan as sole carbon sources. Strains were pure-cultured on minimal seawater agar containing 0.4% glucose and 0.2% casamino acids (w/v). 16S rRNA gene fragments were Sanger-sequenced using primers 27F/907RM at LGC Genomics (Berlin, Germany). Using Geneious Prime, we identified five isolates that match Kongsfjorden ASVs at 100% rRNA gene identity (Supplementary Table S3). These strains were whole-genome sequenced on Sequel IIe and Revio systems (Pacific Biosciences, Menlo Park, CA) taking one 30 h movie per SMRT cell. For this, genomic DNA was extracted using the MasterPureTM Gram Positive DNA Purification Kit (Epicentre Biotechnologies, Germany) according to the manufactureŕs instructions. Samples were pooled equimolar. A template library was prepared using the SMRTbell^®^ prep kit 3.0, following the protocol ‘Preparing whole genome and metagenome libraries’ to ensure optimal primer annealing and polymerase binding to purified SMRTbell template. Loading was done using the calculator in SMRT Link v13.1. Long-read assemblies were performed with the ‘Microbial Genome Analysis’ protocol in SMRT Link v13.1. For potentially mixed cultures (NYA28a, NYA30) the dominant strain was retrieved though an additional assembly with Flye v2.9 [87]. Assemblies was circularized, the start point adjusted (*dnaA* for chromosomes), and corrected via mapping of PacBio reads using Minimap2 v2.28 [88] and variant detection using VarScan v2.28 [89]. Closed genomes were annotated using Bakta v1.11.3 [90]. CAZymes, biosynthetic gene clusters, and metabolic genes were annotated using dbCAN3, antiSMASH v8.0, KofamKOALA, and KEGG Mapper [91–94].

### Carbohydrate quantification

To quantify carbohydrates in particulate (pCCHO, >0.2 μm) and dissolved (dCCHO/dFCHO, <0.2 μm) phases, seawater samples were analyzed according to Zeppenfeld et al. [95, 96] via high-performance anion-exchange chromatography with pulsed amperometric detection (HPAEC-PAD), utilizing a Dionex CarboPac PA20 analytical column (3mm × 150 mm) and a PA20 guard column (3mm × 30 mm). The monosaccharides fucose (Fuc), rhamnose (Rha), arabinose (Ara), galactose (Gal), glucose (Glc), xylose (Xyl), mannose (Man), fructose (Fru), galactosamine (GalN), glucosamine (GlcN), muramic acid (MurAc), galacturonic acid (GalAc), and glucuronic acid (GlcAc) were identified by their retention times. dFCHO comprises the identifiable monosaccharides, whereas dCCHO and pCCHO include those released following acid hydrolysis (0.8 M HCl, 100°C, 20 h). Total combined carbohydrates (CCHO) include both dCCHO and pCCHO, whereas CHO represents the sum of CCHO and dFCHO. Carbohydrates in aerosol particles were measured from aqueous extracts with (CCHO_aer_) or without (dFCHO_aer_) acid hydrolysis.

### Meteorological data and airmass history

Data from the AWIPEV atmospheric observatory (https://doi.org/10.1594/PANGAEA.914979) were used to calculate wind speed, wind direction, air temperature, and air pressure across the aerosol sampling. 48-hour back-trajectories (generated hourly at arrival altitudes of 50, 250, and 1000 m) were computed using the NOAA HYSPLIT model [97] based on GDAS1 meteorological fields (Global Data Assimilation System; 1° latitude/longitude; 3-hourly). Sea-ice concentrations were retrieved from ERDDAP (https://opendap.co-ops.nos.noaa.gov/erddap/index.html). To analyze air mass histories in relation to surface types, back trajectories and sea-ice maps were integrated to calculate the relative residence time of air masses over specific surface areas before sampling. Surface areas were classified into open ocean (sea-ice concentration <15%), marginal ice zone (sea-ice concentration 15−80%), pack ice (sea-ice concentration 80−100%), and land.

### Statistical evaluation

Data was analyzed in RStudio using R v4.5.1 and packages *tidyverse*, *gtools*, *cowplot*, *fuzzyjoin*, *ampvis2*, *vegan*, *UpSetR*, *mixOmics*, *amap*, *ape*, *lubridate*, *FEAST*, *iNEXT*, *rstatix*, *openair*, *ggOceanMaps*, *ggspatial*, *solartime, oce*, *ocedata*, *ncdf4*, *openair*, *scales*, *lubridate*, *cmocean*, *maps*, *mapdata*, *sp*, *rgdal*, *raster*, and *RColorBrewer* [98–125]. ENVO terms were assigned in accordance with Lang-Yona and colleagues [126]. Figures were arranged and beautified using Inkscape. All bioinformatic code and data files are available under https://github.com/matthiaswietz/BelugaMicrobes.

## Data availability

Amplicon sequences are available under ENA accession number PRJEB91661; deposited via the German Federation for Biological Data (GFBio) in compliance with MIxS standards. Closed bacterial genomes are available under ENA accession number PRJEB91655. Carbohydrate and ion concentrations in ULW and SML are available under https://doi.org/10.1594/PANGAEA.982606. Carbohydrate concentrations in aerosol particles are available under https://doi.org/10.1594/PANGAEA.982703.

## ACKNOWLEDGMENTS

Our deepest gratitude to the BELUGA team for their dedication, support, and expertise: Holger Siebert and Birgit Wehner led the field campaigns in 2021 and 2022, with vital contributions by Thomas Conrath, Mona Kellermann, Christian Pilz, Joshua Müller, Michael Lonardi, Jonas Schaefer, Pablo Saavedra Garfias, André Ehrlich, Jens Voigtländer, Elisa Akansu, Jakob Thoböll, and René Rabe. We are indebted to the AWIPEV Observatory, in particular Greg Tran, Fieke Rader, Yohann Dulong and Isabell Schulz, for their immense and friendly support. We thank Kings Bay for creating a professional and pleasant environment. DNA extraction and amplicon sequencing was expertly done by Jakob Barz, Kerstin Korte, and Katja Metfies. We thank Tabea Platz for 16S rRNA sequencing of isolates. We appreciate funding by the German Research Foundation (project number 268020496) within the Transregional Collaborative Research Center TRR 172 “ArctiC Amplification: Climate Relevant Atmospheric and SurfaCe Processes, and Feedback Mechanisms (AC)3” in subprojects B04 and A02. MW was supported by the German Research Foundation priority program SPP 1158 “Antarctic Research with comparative investigations in Arctic ice areas” (project number 522416631).

## SUPPLEMENTARY DATA

**Supplementary Figure S1: ASVs and matching bacterial isolates.** Average relative abundances of ASVs with 100% rRNA identity to genome-sequenced isolates by niche and season.

**Supplementary Figure S2: Fog water microbiomes.** Predominant prokaryotes in fog water, collected in autumn 2021 at the Gruvebadet Observatory, in comparison to ULW and SML.

**Supplementary Figure S3: Carbohydrate concentrations across niches.** Values are µg L^-1^ for ULW and SML, and ng m^-3^ for aerosol particles.

**Supplementary Figure S4: Seasonal meteorological conditions at the AWIPEV Observatory. A:** Wind speed, air temperature, and air pressure (10 meters above ground). **B:** Summary of wind directions and speed.

**Supplementary Figure S5: Wind trajectories.** Animated back-tracking of wind trajectories in autumn 2021 (left) and spring 2022 (right), determined hourly at three arrival altitudes (red 50 m, violet 250 m, pink 1000m).

**Supplementary Table S1:** Sample overview and metadata.

**Supplementary Table S2:** Carbohydrate concentrations and meteorological conditions.

**Supplementary Table S3:** sPLS cluster assignment and genomic traits.

## REFERENCES

1. Wilson B et al. Changes in Marine Prokaryote Composition with Season and Depth Over an Arctic Polar Year. Frontiers in Marine Science 2017;4:95.

2. Wietz M et al. The polar night shift: seasonal dynamics and drivers of Arctic Ocean microbiomes revealed by autonomous sampling. ISME Commun 2021;1:76. 10.1038/s43705-021-00074-4

3. von Jackowski A et al. Variations of microbial communities and substrate regimes in the eastern Fram Strait between summer and fall. Environ Microbiol 2022;24:4124–4136. 10.1111/1462-2920.16036

4. Wietz M et al. The Arctic summer microbiome across Fram Strait: Depth, longitude, and substrate concentrations structure microbial diversity in the euphotic zone. Environ Microbiol 2024;6:e16568. 10.1111/1462-2920.16568

5. Freud E et al. Pan-Arctic aerosol number size distributions: seasonality and transport patterns. Atmospheric Chemistry and Physics 2017;17:8101–8128. 10.5194/acp-17-8101-2017

6. Jang S et al. Large seasonal and interannual variations of biogenic sulfur compounds in the Arctic atmosphere (Svalbard; 78.9°N, 11.9°E). Atmospheric Chemistry and Physics 2021;21:9761–9777. 10.5194/acp-21-9761-2021

7. Jang E et al. Seasonal dynamics of airborne biomolecules influence the size distribution of Arctic aerosols. Environmental Science and Ecotechnology 2024;22:100458. 10.1016/j.ese.2024.100458

8. O’Dowd CD et al. Biogenically driven organic contribution to marine aerosol. Nature 2004;431:676–680. 10.1038/nature02959

9. Quinn PK et al. Chemistry and related properties of freshly emitted sea spray aerosol. Chem Rev 2015;115:4383–4399. 10.1021/cr500713g

10. Šantl-Temkiv T et al. Microbial ecology of the atmosphere. FEMS Microbiol Rev 2022;46:fuac009. 10.1093/femsre/fuac009

11. Aller JY et al. The sea surface microlayer as a source of viral and bacterial enrichment in marine aerosols. Journal of Aerosol Science 2005;36:801–812. 10.1016/j.jaerosci.2004.10.012

12. Orellana MV et al. Marine microgels as a source of cloud condensation nuclei in the high Arctic. Proceedings of the National Academy of Sciences USA 2011;108:13612–13617. 10.1073/pnas.1102457108

13. Cunliffe M et al. Sea surface microlayers: A unified physicochemical and biological perspective of the air–ocean interface. Progress in Oceanography 2013;109:104–116. 10.1016/j.pocean.2012.08.004

14. Quinn PK et al. Contribution of sea surface carbon pool to organic matter enrichment in sea spray aerosol. Nature Geosci 2014;7:228–232. 10.1038/ngeo2092

15. Fahlgren C et al. Seawater mesocosm experiments in the Arctic uncover differential transfer of marine bacteria to aerosols. Environ Microbiol Rep 2015;7:460–470. 10.1111/1758-2229.12273

16. Michaud JM et al. Taxon-specific aerosolization of bacteria and viruses in an experimental ocean-atmosphere mesocosm. Nat Commun 2018;9:2017. 10.1038/s41467-018-04409-z

17. Tignat-Perrier R et al. Global airborne microbial communities controlled by surrounding landscapes and wind conditions. Sci Rep 2019;9:14441. 10.1038/s41598-019-51073-4

18. Tignat-Perrier R et al. Seasonal shift in airborne microbial communities. Sci Tot Environ 2020;716:137129. 10.1016/j.scitotenv.2020.137129

19. Uetake J et al. Airborne bacteria confirm the pristine nature of the Southern Ocean boundary layer. Proceedings of the National Academy of Sciences USA 2020;117:13275–13282. 10.1073/pnas.2000134117

20. Archer S et al. Global biogeography of atmospheric microorganisms reflects diverse recruitment and environmental filtering. 2022. Preprint from Research Square, 2022.

21. Zeppenfeld S et al. Marine carbohydrates in Arctic aerosol particles and fog – diversity of oceanic sources and atmospheric transformations. Atmospheric Chemistry and Physics 2023;23:15561–15587. 10.5194/acp-23-15561-2023

22. Madawala CK et al. Effects of wind speed on size-dependent morphology and composition of sea spray aerosols. ACS Earth Space Chem 2024;8:1609–1622. 10.1021/acsearthspacechem.4c00119

23. Honeyman AS, Day ML, Spear JR. Regional fresh snowfall microbiology and chemistry are driven by geography in storm-tracked events, Colorado, USA. PeerJ 2018;6:e5961. 10.7717/peerj.5961

24. Park J et al. Shipborne observations reveal contrasting Arctic marine, Arctic terrestrial and Pacific marine aerosol properties. Atmospheric Chemistry and Physics 2020;20:5573–5590. 10.5194/acp-20-5573-2020

25. Steiner AL. Role of the terrestrial biosphere in atmospheric chemistry and climate. Acc Chem Res 2020;53:1260–1268. 10.1021/acs.accounts.0c00116

26. Choufany M et al. Inferring long-distance connectivity shaped by air-mass movement for improved experimental design in aerobiology. Sci Rep 2021;11:11093. 10.1038/s41598-021-90733-2

27. Feltracco M et al. Free and combined L- and D-amino acids in Arctic aerosol. Chemosphere 2019;220:412–421. 10.1016/j.chemosphere.2018.12.147

28. Castenschiold CDF et al. Atmospheric Biogenic Ice-Nucleating Particles Link to Microbial Communities in the Arctic Marine Environment in Western Greenland. Environ Sci Technol 2025. 10.1021/acs.est.5c03650

29. Schneider SR et al. Abiotic emission of volatile organic compounds from the ocean surface: relationship to seawater composition. ACS Earth Space Chem 2024;9:1913–1923. 10.1021/acsearthspacechem.4c00163

30. Feltracco M et al. Airborne bacteria and particulate chemistry capture phytoplankton bloom dynamics in an Arctic fjord. Atmospheric Environment 2021;256:118458. 10.1016/j.atmosenv.2021.118458

31. Matulová M et al. Biotransformation of various saccharides and production of exopolymeric substances by cloud-borne *Bacillus* sp. 3B6. Environ Sci Technol 2014;48:14238–14247. 10.1021/es501350s

32. Hegseth EN, et al. Phytoplankton seasonal dynamics in Kongsfjorden, Svalbard and the adjacent shelf. In: Hop H, Wiencke C (eds), The Ecosystem of Kongsfjorden, Svalbard. Cham: Springer International Publishing, 2019, 173–227.

33. Tverberg V, et al. The Kongsfjorden Transect: seasonal and inter-annual variability in hydrography. In: Hop H, Wiencke C (eds), The Ecosystem of Kongsfjorden, Svalbard. Cham: Springer International Publishing, 2019, 49–104.

34. Herbert LC et al. Tight benthic-pelagic coupling drives seasonal and interannual changes in iron-sulfur cycling in Arctic fjord sediments (Kongsfjorden, Svalbard). Journal of Marine Systems 2021;225:103645. 10.1016/j.jmarsys.2021.103645

35. Jain A et al. Spatially resolved assembly, connectivity and structure of particle-associated and free-living bacterial communities in a high Arctic fjord. FEMS Microbiol Ecol 2021;97:fiab139. 10.1093/femsec/fiab139

36. Assmy P et al. Plankton dynamics in Kongsfjorden during two years of contrasting environmental conditions. Progress in Oceanography 2023;213:102996. 10.1016/j.pocean.2023.102996

37. Hoppe CJM et al. Photosynthetic light requirement near the theoretical minimum detected in Arctic microalgae. Nat Commun 2024;15:7385. 10.1038/s41467-024-51636-8

38. Hoppe CJM et al. The effects of biomass depth distribution on phytoplankton spring bloom dynamics and composition in an Arctic fjord. Elementa: Science of the Anthropocene 2024;12:00137. 10.1525/elementa.2023.00137

39. Teeling H et al. Substrate-controlled succession of marine bacterioplankton populations induced by a phytoplankton bloom. Science 2012;336:608–611. 10.1126/science.1218344

40. Jain A et al. Response of bacterial communities from Kongsfjorden (Svalbard, Arctic Ocean) to macroalgal polysaccharide amendments. Mar Environ Res 2020;155:104874. 10.1016/j.marenvres.2020.104874

41. Zeppenfeld S et al. Marine Carbohydrates and Other Sea Spray Aerosol Constituents Across Altitudes in the Lower Troposphere of Ny-Ålesund, Svalbard. EGUsphere 2025;1–49. 10.5194/egusphere-2025-4336

42. Priest T et al. Atlantic water influx and sea-ice cover drive taxonomic and functional shifts in Arctic marine bacterial communities. ISME J 2023;17:1612–1625. 10.1038/s41396-023-01461-6

43. Priest T et al. Seasonal recurrence and modular assembly of an Arctic pelagic marine microbiome. Nat Commun 2025;16:1326. 10.1038/s41467-025-56203-3

44. Ontiveros VJ et al. General decline in the diversity of the airborne microbiota under future climatic scenarios. Sci Rep 2021;11:20223. 10.1038/s41598-021-99223-x

45. Delpech L-M et al. Terrestrial inputs shape coastal bacterial and archaeal communities in a high Arctic fjord (Isfjorden, Svalbard). Front Microbiol 2021;12:614634. 10.3389/fmicb.2021.614634

46. Alsante AN, Thornton DCO, Brooks SD. Ocean aerobiology. Front Microbiol 2021;12:764178. 10.3389/fmicb.2021.764178

47. Francis B et al. North Sea spring bloom-associated Gammaproteobacteria fill diverse heterotrophic niches. Environ Microbiome 2021;16:15. 10.1186/s40793-021-00385-y

48. Silva AN et al. Meta-analytical insights into organic matter enrichment in the surface microlayer. EGUsphere 2025;1–30. 10.5194/egusphere-2025-4050

49. Giovannoni SJ. SAR11 bacteria: the most abundant plankton in the oceans. Annu Rev Mar Sci 2017;9:231–255. 10.1146/annurev-marine-010814-015934

50. Wang F-Q et al. Particle-attached bacteria act as gatekeepers in the decomposition of complex phytoplankton polysaccharides. Microbiome 2024;12:32. 10.1186/s40168-024-01757-5

51. Yu L. On sea surface salinity skin effect induced by evaporation and implications for remote sensing of ocean salinity. Journal of Physical Oceanography 2010;40:85–102. 10.1175/2009JPO4168.1

52. Santos AL et al. Diversity in UV sensitivity and recovery potential among bacterioneuston and bacterioplankton isolates. Lett Appl Microbiol 2011;52:360–366. 10.1111/j.1472-765X.2011.03011.x

53. Sedkova N et al. Diversity of carotenoid synthesis gene clusters from environmental Enterobacteriaceae strains. Appl Environ Microbiol 2005;71:8141–8146. 10.1128/AEM.71.12.8141-8146.2005

54. Dieser M, Greenwood, Mark, and Foreman CM. Carotenoid pigmentation in Antarctic heterotrophic bacteria as a strategy to withstand environmental stresses. Arctic, Antarctic, and Alpine Research 2010;42:396–405. 10.1657/1938-4246-42.4.396

55. Vila E et al. Carotenoids from heterotrophic bacteria isolated from Fildes Peninsula, King George Island, Antarctica. Biotechnology Reports 2019;21:e00306. 10.1016/j.btre.2019.e00306

56. Coppola D et al. Biodiversity of UV-resistant bacteria in Antarctic aquatic environments. Journal of Marine Science and Engineering 2023;11:968. 10.3390/jmse11050968

57. Wurl O, Miller L, Vagle S. Production and fate of transparent exopolymer particles in the ocean. Journal of Geophysical Research: Oceans 2011;116. 10.1029/2011JC007342

58. Campbell BJ et al. The effect of nutrient deposition on bacterial communities in Arctic tundra soil. Environmental Microbiology 2010;12:1842–1854. 10.1111/j.1462-2920.2010.02189.x

59. Zhang D-C et al. *Arthrobacter alpinus* sp. nov., a psychrophilic bacterium isolated from alpine soil. Int J Syst Evol Microbiol 2010;60:2149–2153. 10.1099/ijs.0.017178-0

60. Keuschnig C et al. Selection processes of Arctic seasonal glacier snowpack bacterial communities. Microbiome 2023;11:35. 10.1186/s40168-023-01473-6

61. Jones AM, Harrison RM. The effects of meteorological factors on atmospheric bioaerosol concentrations--a review. Sci Total Environ 2004;326:151–180. 10.1016/j.scitotenv.2003.11.021

62. Qi J et al. Estimation of dry deposition fluxes of particulate species to the water surface in the Qingdao area, using a model and surrogate surfaces. Atmospheric Environment 2005;39:2081–2088. 10.1016/j.atmosenv.2004.12.017

63. Jensen LZ et al. Seasonal Variation of the Atmospheric Bacterial Community in the Greenlandic High Arctic Is Influenced by Weather Events and Local and Distant Sources. Frontiers in Microbiology 2022;13:909980. 10.3389/fmicb.2022.909980

64. Malard LA et al. Snow microorganisms colonise Arctic soils following snow melt. Microb Ecol 2023;86:1661–1675. 10.1007/s00248-023-02204-y

65. Vijayakumar VR, Dharumadurai D. Chapter 10 - Endolichenic actinobacterial association in fruticose, foliose, and crustose lichens. In: Dharumadurai D (ed.), Microbial Symbionts. Academic Press, 2023, 177–191.

66. Ruben M et al. Microbial communities degrade ancient permafrost-derived organic matter in Arctic seawater. Journal of Geophysical Research: Biogeosciences 2024;129:e2023JG007936. 10.1029/2023JG007936

67. Bhaskar JT et al. Variation of phytoplankton assemblages of Kongsfjorden in early autumn 2012: a microscopic and pigment ratio-based assessment. Environ Monit Assess 2016;188:224. 10.1007/s10661-016-5220-8

68. Zeng Y-X et al. Diversity of bacterial dimethylsulfoniopropionate degradation genes in surface seawater of Arctic Kongsfjorden. Scientific Reports 2016;6:33031. 10.1038/srep33031

69. Belevich TA, Milyutina IA, Troitsky AV. Seasonal variability of photosynthetic microbial eukaryotes (<3 µm) in the Kara Sea revealed by 18S rDNA metabarcoding of sediment trap fluxes. Plants 2021;10:2394. 10.3390/plants10112394

70. Terrado R et al. Protist community composition during spring in an Arctic flaw lead polynya. Polar Biol 2011;34:1901–1914. 10.1007/s00300-011-1039-5

71. Meshram AR et al. Microbial eukaryotes in an Arctic under-ice spring bloom north of Svalbard. Front Microbiol 2017;8:1099. 10.3389/fmicb.2017.01099

72. Creamean JM et al. Ice nucleating particles carried from below a phytoplankton bloom to the Arctic atmosphere. Geophysical Research Letters 2019;46:8572–8581. 10.1029/2019GL083039

73. Marks R et al. Rising bubbles as mechanism for scavenging and aerosolization of diatoms. Journal of Aerosol Science 2019;128:79–88. 10.1016/j.jaerosci.2018.12.003

74. Düsedau L et al. Kelp forest community structure and demography in Kongsfjorden (Svalbard) across 25 years of Arctic warming. Ecology and Evolution 2024;14:e11606. 10.1002/ece3.11606

75. Sinnett G et al. Contribution of large marine aerosols in phytoplankton dispersal. Environ Sci Technol 2025;59:7348–7356. 10.1021/acs.est.4c14473

76. Wang Y et al. A Novel Cold-Adapted Nitronate Monooxygenase from Psychrobacter sp. ANT206: Identification, Characterization and Degradation of 2-Nitropropane at Low Temperature. Microorganisms 2024;12:2100. 10.3390/microorganisms12102100

77. Sadler MC et al. Genome restructuring and adaptation in Arctic marine bacteria. mBio. 2025. 2025., 16: e01555-25

78. Jaarsma AH et al. The undiscovered biosynthetic potential of the Greenland Ice Sheet microbiome. Front Microbiol 2023;14:1285791. 10.3389/fmicb.2023.1285791

79. Barberán A et al. Structure, inter-annual recurrence, and global-scale connectivity of airborne microbial communities. Sci Total Environ 2014;487:187–195. 10.1016/j.scitotenv.2014.04.030

80. Zeppenfeld S et al. Glucose as a potential chemical marker for ice nucleating activity in Arctic seawater and melt pond samples. Environ Sci Technol 2019;53:8747–8756. 10.1021/acs.est.9b01469

81. Parada AE, Needham DM, Fuhrman JA. Every base matters: assessing small subunit rRNA primers for marine microbiomes with mock communities, time series and global field samples. Environ Microbiol 2016;18:1403–1414. 10.1111/1462-2920.13023

82. Fadeev E et al. Microbial communities in the east and west Fram Strait during sea ice melting season. Front Mar Sci 2018;5:429. 10.3389/fmars.2018.00429

83. Callahan BJ et al. DADA2: High-resolution sample inference from Illumina amplicon data. Nat Methods 2016;13:581–583. 10.1038/nmeth.3869

84. Quast C et al. The SILVA ribosomal RNA gene database project: improved data processing and web-based tools. Nucleic Acids Res 2013;41:D590–596. 10.1093/nar/gks1219

85. Guillou L et al. The Protist Ribosomal Reference database (PR2): a catalog of unicellular eukaryote Small Sub-Unit rRNA sequences with curated taxonomy. Nucleic Acids Res 2013;41:D597–D604. 10.1093/nar/gks1160

86. Zech H et al. Growth phase-dependent global protein and metabolite profiles of *Phaeobacter gallaeciensis* strain DSM 17395, a member of the marine *Roseobacter* clade. Proteomics 2009;9:3677–3697. 10.1002/pmic.200900120

87. Kolmogorov M et al. Assembly of long, error-prone reads using repeat graphs. Nat Biotechnol 2019;37:540–546. 10.1038/s41587-019-0072-8

88. Li H. New strategies to improve minimap2 alignment accuracy. Bioinformatics 2021;37:4572–4574. 10.1093/bioinformatics/btab705

89. Koboldt DC et al. VarScan 2: somatic mutation and copy number alteration discovery in cancer by exome sequencing. Genome Res 2012;22:568–76. 10.1101/gr.129684.111

90. Schwengers O et al. Bakta: rapid and standardized annotation of bacterial genomes via alignment-free sequence identification. Microbial Genomics 2021;7:000685. 10.1099/mgen.0.000685

91. Zheng J, et al. dbCAN3: automated carbohydrate-active enzyme and substrate annotation. Nucleic Acids Research 2023;51:W115–W121. 10.1093/nar/gkad328

92. Blin K et al. antiSMASH 8.0: extended gene cluster detection capabilities and analyses of chemistry, enzymology, and regulation. Nucleic Acids Research 2025;53:W32–W38. 10.1093/nar/gkaf334

93. Aramaki T et al. KofamKOALA: KEGG Ortholog assignment based on profile HMM and adaptive score threshold. Bioinformatics 2020;36:2251–2252. 10.1093/bioinformatics/btz859

94. Kanehisa M, Sato Y, Kawashima M. KEGG mapping tools for uncovering hidden features in biological data. Protein Sci 2022;31:47–53. 10.1002/pro.4172

95. Zeppenfeld S et al. A protocol for quantifying mono- and polysaccharides in seawater and related saline matrices by electro-dialysis (ED) – combined with HPAEC-PAD. Ocean Science 2020;16:817–830. 10.5194/os-16-817-2020

96. Zeppenfeld S et al. Aerosol marine primary carbohydrates and atmospheric transformation in the Western Antarctic Peninsula. ACS Earth Space Chem 2021;5:1032–1047. 10.1021/acsearthspacechem.0c00351

97. Stein AF et al. NOAA’s HYSPLIT Atmospheric Transport and Dispersion Modeling System. Bulletin of the American Meteorological Society 2015;96:2059–2077. 10.1175/BAMS-D-14-00110.1

98. Carslaw DC, Ropkins K. openair — An R package for air quality data analysis. 2012. 2012.

99. Oksanen J et al. Vegan: community ecology package. 2013. 2013.

100. Hsieh TC, Ma KH, Chao A. iNEXT: an R package for rarefaction and extrapolation of species diversity (Hill numbers). Methods Ecol Evol 2016;7:1451–1456. 10.1111/2041-210X.12613

101. Conway J, Lex A, Gehlenborg N. UpSetR: an R package for the visualization of intersecting sets and their properties. Bioinformatics. 2017. 2017., 33: 2938–2940

102. Rohart F et al. mixOmics: An R package for ‘omics feature selection and multiple data integration. PLOS Computational Biology 2017;13:e1005752. 10.1371/journal.pcbi.1005752

103. Andersen KSS et al. ampvis2: an R package to analyse and visualise 16S rRNA amplicon data. bioRxiv 2018;299537. 10.1101/299537

104. Paradis E, Schliep K. ape 5.0: an environment for modern phylogenetics and evolutionary analyses in R. Bioinformatics 2019;35:526–528. 10.1093/bioinformatics/bty633

105. Shenhav L et al. FEAST: fast expectation-maximization for microbial source tracking. Nat Methods 2019;16:627–632. 10.1038/s41592-019-0431-x

106. Wickham H et al. Welcome to the Tidyverse. J Open Source Softw 2019;4:1686. 10.21105/joss.01686

107. Robinson D. fuzzyjoin: join tables together on inexact matching. 2020. 2020.

108. Warnes G, Bolker B, Lumley T. gtools package. 2020. 2020.

109. Dunnington D. ggspatial: spatial data framework for ggplot2. 2023. 2023.

110. Kassambara A. rstatix: pipe-friendly framework for basic statistical tests. 2023. 2023.

111. Lucas A. amap: Another Multidimensional Analysis Package. 2024. 2024.

112. Vihtakari M. ggOceanMaps: plot data on oceanographic maps using ‘ggplot2’. 2024. 2024.

113. Wilke CO. cowplot: streamlined plot theme and plot annotations for ‘ggplot2’. 2024. 2024.

114. Wutzler T. solartime: utilities dealing with solar time such as sun position and time of sunrise. 2024. 2024.

115. Grolemund G, Wickham H. Dates and times made easy with lubridate. 2011. 2011.

116. Bivand RS, Pebesma E, Gomez-Rubio V. Applied spatial data analysis with R, Second edition. 2013. Springer, NY, 2013.

117. Thyng KM, et al. True colors of oceanography: Guidelines for effective and accurate colormap selection. 2016. 2016.

118. Wickham H, Pedersen TL, Seidel D. scales: scale functions for visualization. 2025. 2025.

119. Becker OS code by RA, Brownrigg ARWR version by R. mapdata: Extra Map Databases. 2022. 2022.

120. Kelley D, Richards C. oce: analysis of oceanographic data. 2025. 2025.

121. Kelley D, Richards C. ocedata: oceanographic data sets for ‘oce’ package. 2022. 2022.

122. Neuwirth E. RColorBrewer: ColorBrewer Palettes. 2022. 2022.

123. Becker RA, et al. maps: draw geographical maps. 2025. 2025.

124. Hijmans RJ. raster: geographic data analysis and modeling. 2025. 2025.

125. Pierce D. ncdf4: Interface to Unidata netCDF (Version 4 or Earlier) Format Data Files. 2025. 2025.

126. Lang-Yona N et al. Terrestrial and marine influence on atmospheric bacterial diversity over the north Atlantic and Pacific Oceans. Commun Earth Environ 2022;3:1–10. 10.1038/s43247-022-00441-6

