## Supplementary figures and images for "Seasonal connectivity of microbes and carbohydrates between ocean, atmosphere, and cryosphere in Kongsfjorden (Svalbard, Arctic Ocean)"

### Supplementary Figure S1

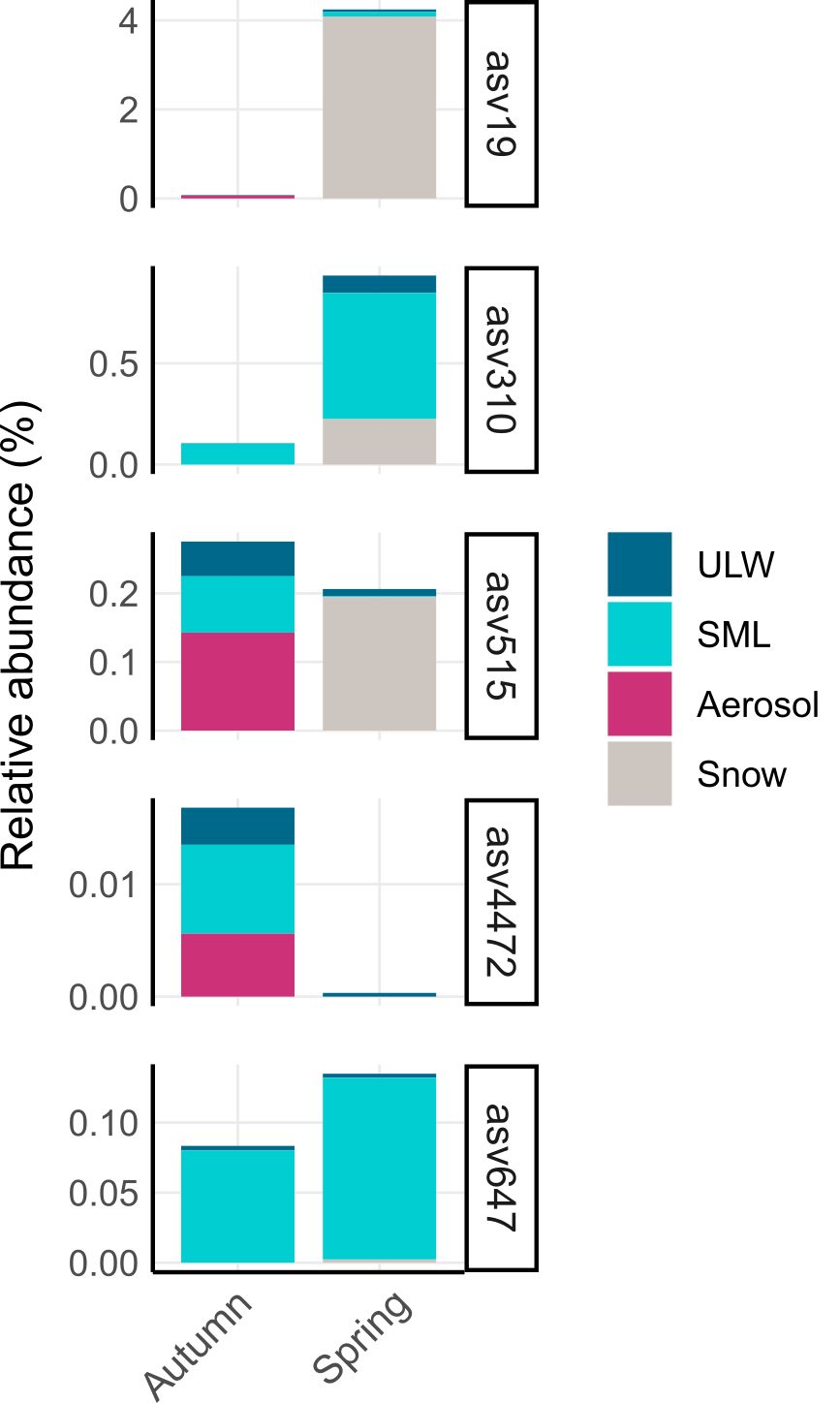

### Supplementary Figure S2

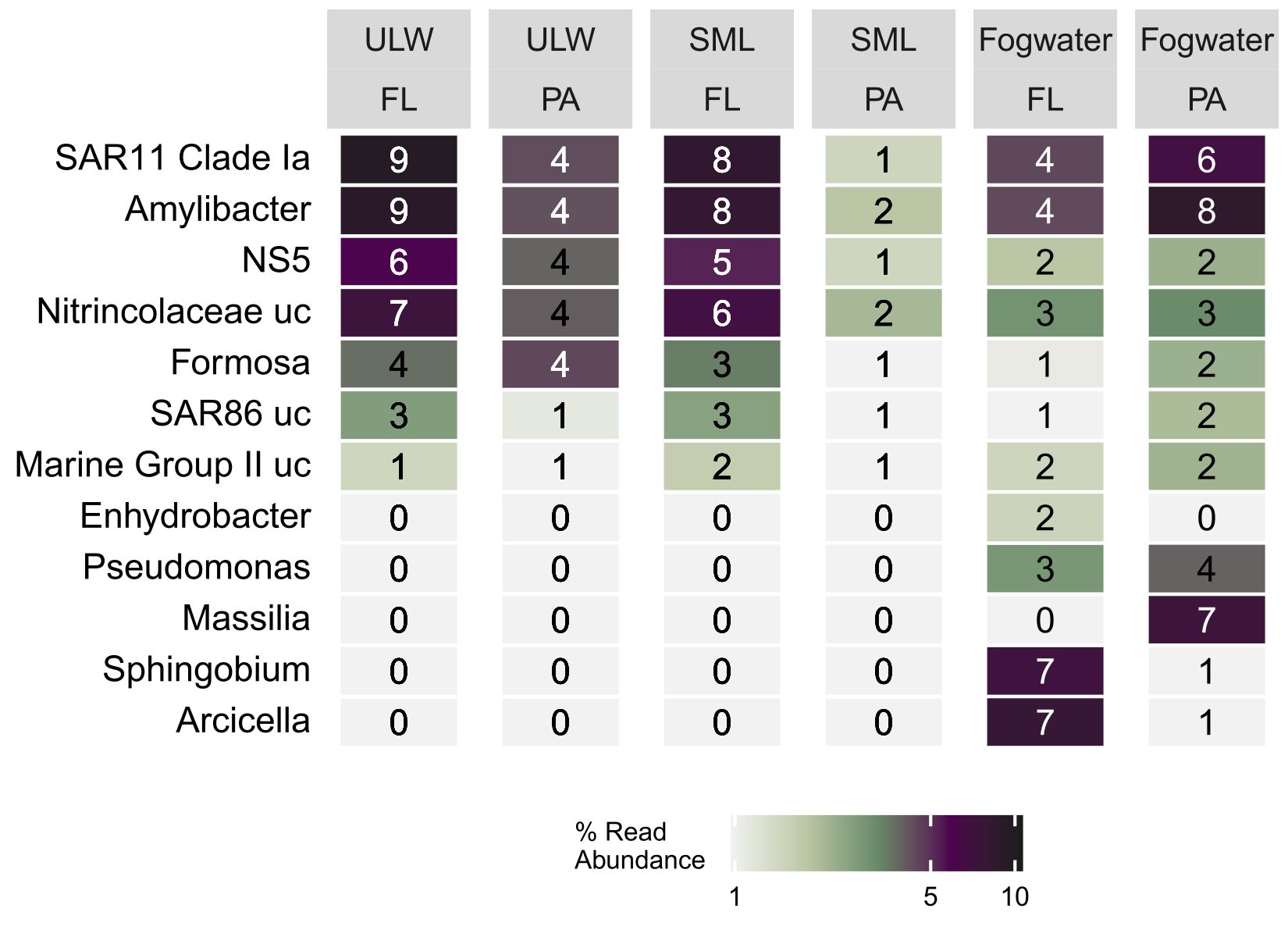

### Supplementary Figure S3

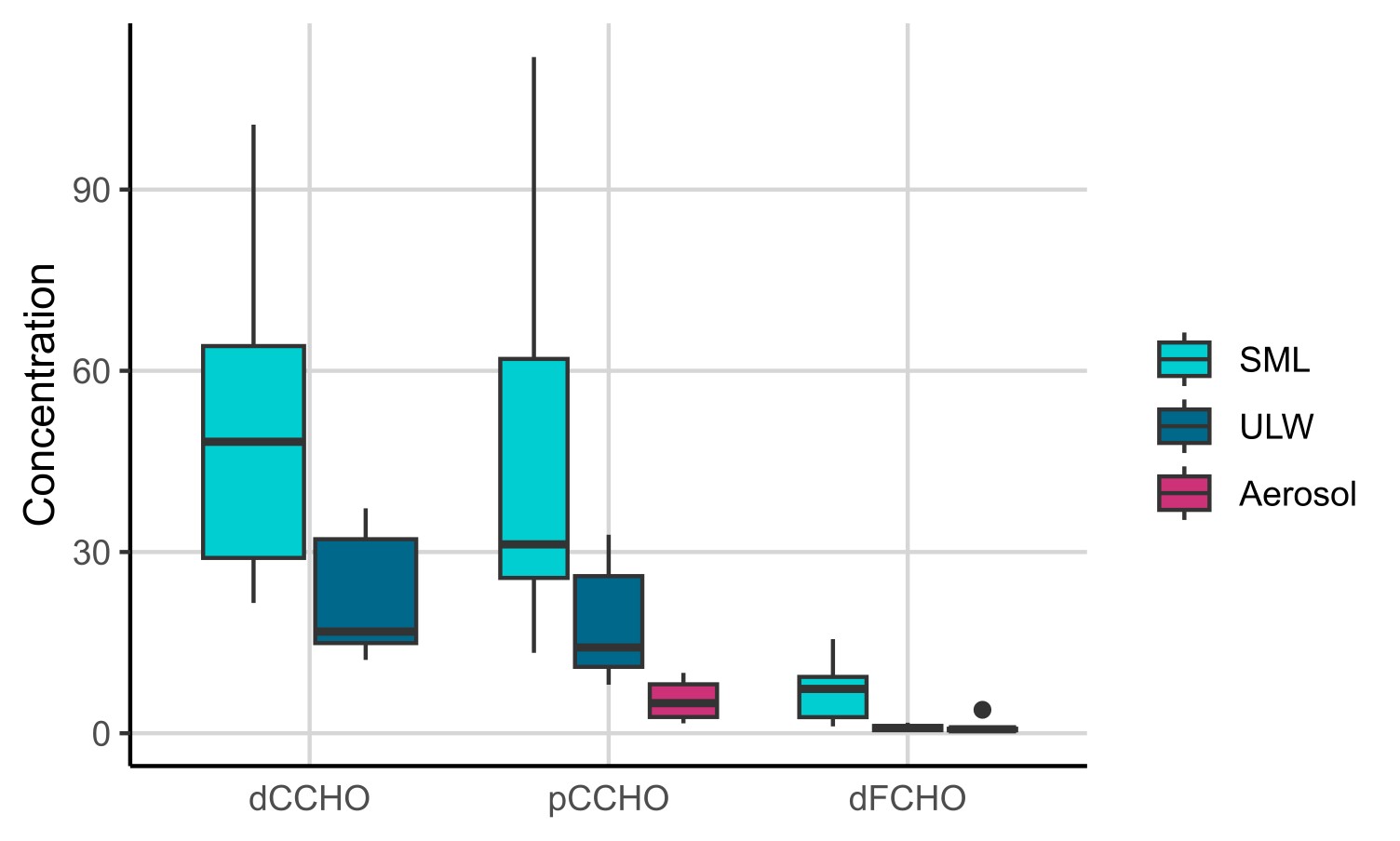

### Supplementary Figure S4

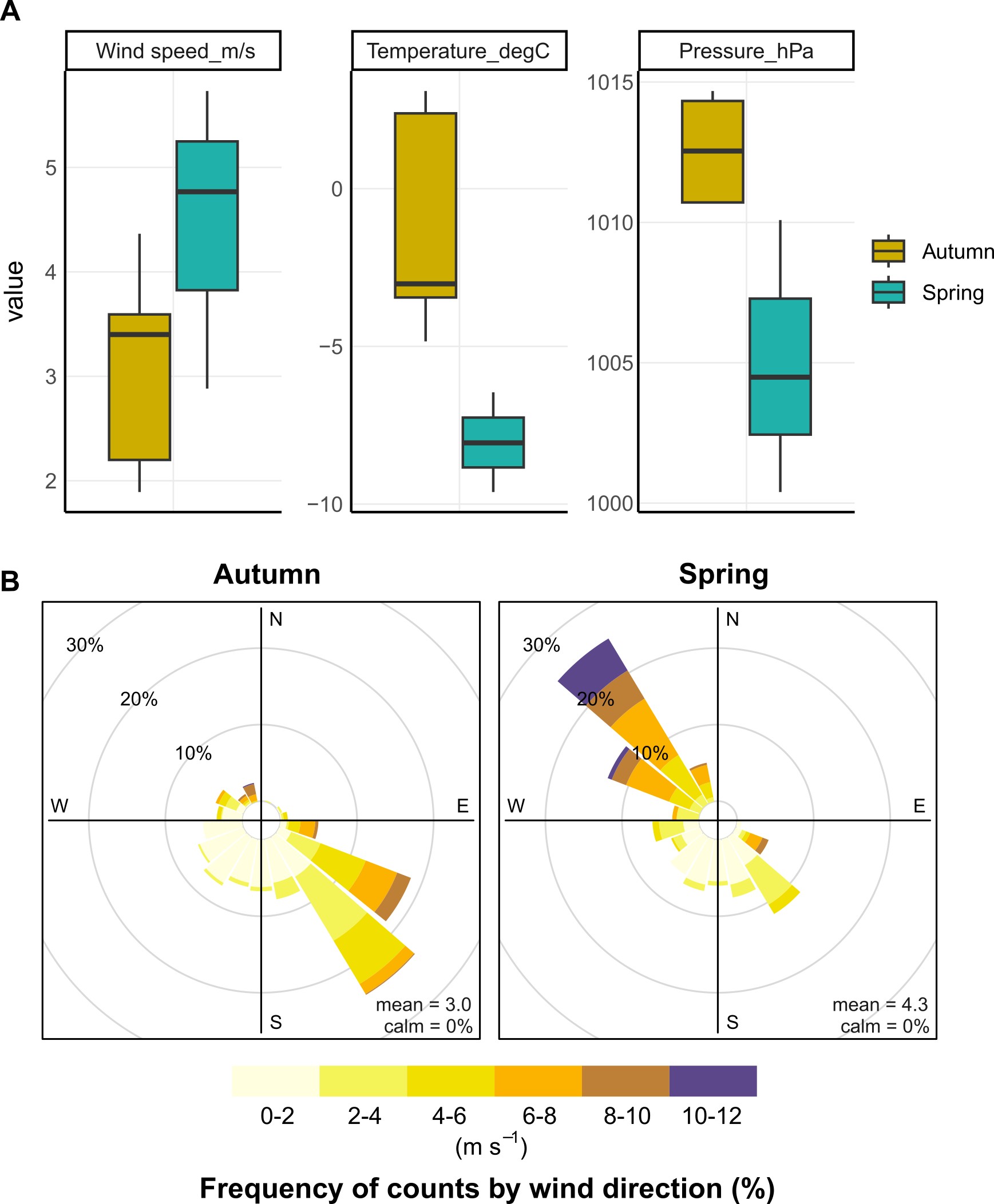

### Supplementary Figure S5

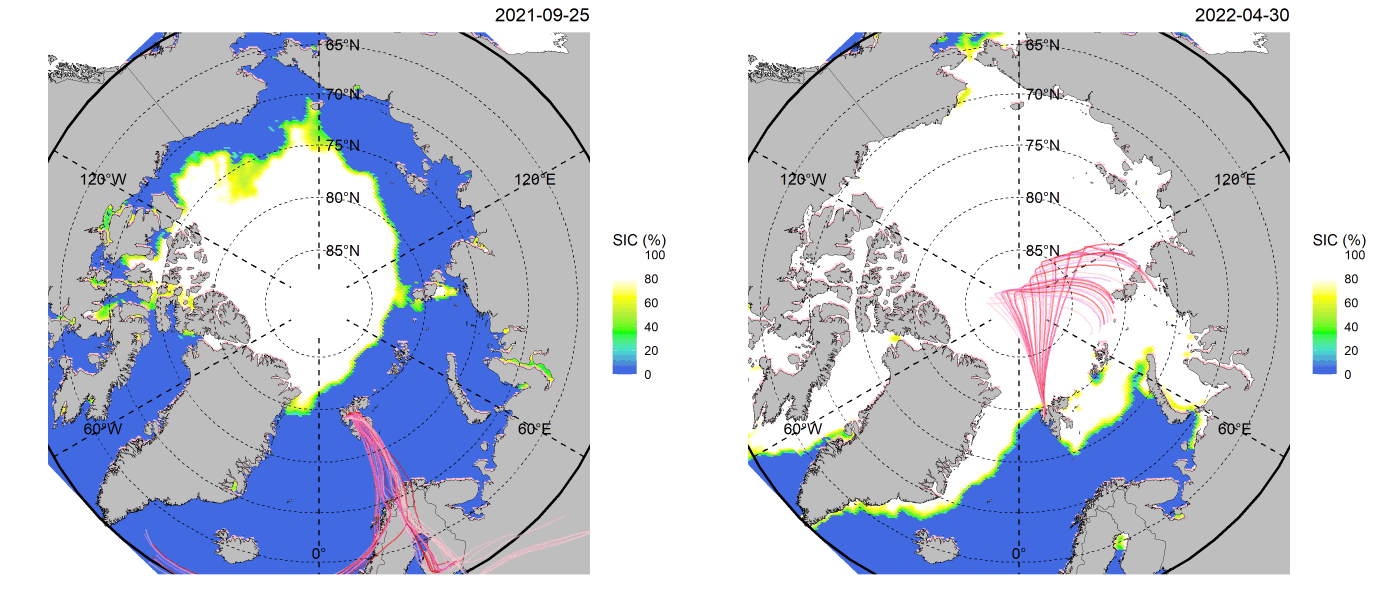
